# Integrative transcriptomic and proteomic meta-analysis of Zika viral infection reveals potential mechanisms for oncolytic therapy in neuroblastoma

**DOI:** 10.1101/2022.11.14.516401

**Authors:** Matt Sherwood, Yilu Zhou, Yi Sui, Yihua Wang, Paul Skipp, Carolini Kaid, Juliet Gray, Keith Okamoto, Rob M. Ewing

## Abstract

**BACKGROUND:** Paediatric neuroblastoma and brain tumours account for a third of all childhood cancer-related mortality. High-risk neuroblastoma is highly aggressive and survival is poor despite intensive multi-modal therapies with significant toxicity. Novel therapies are desperately needed. The Zika virus (ZIKV) is neurotropic and there is growing interest in employing ZIKV as a potential therapy against paediatric nervous system tumours, including neuroblastoma.

**METHODS:** Here, we perform an extensive meta-analysis of ZIKV infection studies to identify molecular mechanisms that may govern the oncolytic response in neuroblastoma cells. We summarise the neuroblastoma cell lines and ZIKV strains utilised and re-evaluate the infection data to deduce the susceptibility of neuroblastoma to the ZIKV oncolytic response. Integrating transcriptomics, interaction proteomics, dependency factor and compound datasets we show the involvement of multiple host systems during ZIKV infection.

**RESULTS:** We identify that most paediatric neuroblastoma cell lines are highly susceptible to ZIKV infection and that the PRVABC59 ZIKV strain is the most promising candidate for neuroblastoma oncolytic virotherapy. ZIKV induces TNF signalling, lipid metabolism, the Unfolded Protein Response (UPR), and downregulates cell cycle and DNA replication processes. ZIKV is dependent on SREBP-regulated lipid metabolism and three protein complexes; V-ATPase, ER Membrane Protein Complex (EMC) and mammalian translocon. We propose ZIKV nonstructural protein 4B (NS4B) as a likely mediator of ZIKVs interaction with IRE1-mediated UPR, lipid metabolism and mammalian translocon.

**CONCLUSIONS:** Our work provides a significant understanding of ZIKV infection in neuroblastoma cells, which will facilitate the progression of ZIKV-based oncolytic virotherapy through pre-clinical research and clinical trials.

**KEYPOINTS:**

- The Zika virus may provide the basis for an oncolytic virotherapy against Neuroblastoma
- Most paediatric neuroblastoma cell lines are susceptible to Zika viral infection
- We identified molecular mechanisms that may induce the oncolytic response in Neuroblastoma

**Contribution to the field:** The ability to both induce direct oncolysis and provoke an anti-tumoral immune response makes oncolytic virotherapy an attractive candidate to combat aggressive and heterogenous cancers, such as high-risk neuroblastoma. To progress oncolytic virotherapy to clinical trial it is essential to understand the host mechanisms the virus manipulates to kill cancer cells, alongside any pathology as a consequence of infection of normal cells. Here, we show that ZIKV efficiently infects and induces oncolysis of paediatric neuroblastoma cells and propose a potential TNF pathway-driven immune response. ZIKV’s specificity for infection of nervous system cancer cells, while rarely causing nervous system-related pathology in young children, addresses many of its safety concerns. The inclusion of more effective and less toxic novel therapies, such as a potential ZIKV-based therapeutic, in multimodal treatment regimens will pave the way for improving patient long-term health and overall survival.

## INTRODUCTION

Neuroblastoma is the most common extracranial solid cancer in children, accounting for 6-10% of all paediatric cancers and disproportionately causing 12-15% of paediatric cancer-related deaths (1). It is an embryonal tumour originating from transformed cells of neural crest lineage and predominately forms tumours in the adrenal medulla and paraspinal sympathetic ganglia. Whilst the majority of patients are diagnosed by the age of 5 years, the median age of patients is 18 months. Prognosis is highly heterogenous and can be predicted by a number of factors including presence of metastatic disease, age, chromosomal aberrations and molecular signatures such as MYC-N amplification (2). Patients are categorised according to internationally agreed risk groups (INRG), and treatment is stratified accordingly.

Outcome for low- and intermediate-risk neuroblastoma is good, with some patients requiring little or no treatment. However, approximately 50% of patients have high-risk disease, for which prognosis is poor, with overall survival of less than 60% (3). Current high-risk neuroblastoma treatment regimens are aggressive. These include multiple rounds of induction chemotherapy, surgical resection, myeloablative chemotherapy, autologous stem cell transplantation and post-consolidation therapy such as immunotherapy (4). The aggressive nature of this regimen carries significant treatment-related mortality and frequently results in long-term toxicities and sequelae impacting the quality of life for surviving patients. Consequently, there is a clear and unmet need for safer and less toxic treatment regimens to combat high-risk neuroblastoma.

Oncolytic virotherapy exploits viruses that preferentially infect and destroy cancer cells via two distinct routes of therapeutic action. Following infection, intense viral replication induces oncolysis, releasing virions into the tumour microenvironment to infect neighbouring tumour cells. Induction of a tumour-specific immune response is a crucial secondary mechanism employed by oncolytic virotherapy that can address highly heterogeneous tumours such as high-risk neuroblastoma and CNS tumours. There is significant interest in combining immuno-modulating cancer therapies with oncolytic virotherapy to augment the anti-tumoral immune response. Oncolytic virotherapy clinical studies have in general reported low toxicity and minimal adverse effects in patients, mainly low-grade constitutional symptoms (5).

ZIKV is a mosquito-borne flavivirus consisting of historical African and epidemic-associated Asian lineages. The latter is neurotropic and can cause microcephaly in the developing fetus through infection of neural stem and progenitor cells, causing cell death and growth reduction (6, 7). In contrast, ZIKV rarely causes adverse effects in children and adults, with the majority of cases (50-80%) being asymptomatic (8). In symptomatic children, ZIKV may cause short-term side effects, namely rash, fever and gastrointestinal symptoms, and in rare instances in adults can cause more severe conditions, such as Guillain-Barré Syndrome, meningitis and encephalitis (8, 9).

Since 2017, the concept of employing the ZIKV as oncolytic virotherapy against brain tumours has gained momentum. ZIKV induces an oncolytic event in infected paediatric brain tumour cells *in vitro* and *in vivo* assays and induces an immune response against spontaneous canine brain tumours (10–12). Paediatric neuroblastoma, like paediatric brain tumours, are predominantly tumours of the nervous system consisting of cancerous cells with neural characteristics. An initial study assessing ZIKV infection in multiple neuroblastoma cell lines demonstrated ZIKV’s potential as a novel neuroblastoma oncolytic virotherapy (13). Here, we survey over 35 studies that have used neuroblastoma cell lines to model ZIKV infection. These studies focused on understanding ZIKV pathology and assessing anti-viral compounds. Through re-analysis and integration of the transcriptomics, proteomics and dependency factor screens from these studies, we identify multiple molecular mechanisms implicated in ZIKV infection of neuroblastoma and aim to determine its potential as oncolytic virotherapy.

## METHODS

### RNA-Seq data processing

RNA-Seq data files (.fastq.gz paired-end) of ZIKV-infected paediatric neuroblastoma SH-SY5Y cells were acquired from the European Nucleotide Archive (ENA) (accession: PRJNA630088). Our RNA-Seq processing pipeline consisted of FastQC (V0.11.9-0), Trim Galore (V0.6.6-0), HISAT2 (V2.2.0), Samtools (V1.11) and Subread (V2.0.1). Reads were aligned against the Homo sapiens GRCh38 genome.

### Differential Gene Expression and Pathway Analysis

Differential gene expression analysis was performed using DeSeq2 to compare the triplicate of ZIKV-infected SH-SY5Y cells versus the triplicate of non-infected SH-SY5Y control cells. Differentially expressed genes (DEG) were plotted on bar charts, volcano and scatter plots using GraphPad PRISM (9.2.0). DEGs (padj < 0.05, fold change > 1.5) were submitted to DAVID as official gene symbols for Gene Ontology (Biological Process Direct), KEGG and Reactome pathway analysis. ZIKV-induced DEGs were mapped onto KEGG pathways using Pathview. ZIKV NS4B host interaction partners were also submitted to DAVID to identify the interaction between NS4B and host pathways. Significance values of DEG and pathway analysis were corrected for multiple testing using the Benjamini and Hochberg method (padj < 0.05).

### Interactome and ZIKV Dependency Factor analysis

The ZIKV NS4B interactome in SK-N-BE(2) paediatric neuroblastoma cells were sourced from IMEx (IM-26452) and mapped in Cytoscape (3.9.1). 130 host NS4B interaction partners were submitted to STRING for high confidence (0.7) evidence-based physical subnetwork analysis, followed by integration with the NS4B interactome to identify the interaction of NS4B with host protein complexes. Additional data incorporated into the Cytoscape map includes known ZIKV dependency factors and the interaction of ZIKV NS2B-3 with the Mammalian Translocon. ZIKV dependency factors were sourced for paediatric SK-N-BE(2) neuroblastoma (14), GSC (15), hiPSC-NPC (16), HEK293FT (15) and HeLa cells (17).

## RESULTS

### ZIKV displays strong oncolytic properties against neuroblastoma cells

ZIKV infects and significantly reduces the cell viability of a multitude of neuroblastoma cell lines from both primary tumour and metastatic sites (Table 1). ZIKV can significantly reduce neuroblastoma cell viability using MOI as low as 0.001 (18). The cell viability of 10/15 neuroblastoma cell lines is significantly reduced to below 20% following ZIKV infection and these observations are apparent despite the differences in the cell line, ZIKV strain, viral MOI and the type of assay performed (Table 1). SK-N-Be(1) and SK-N-Be(2) cells are from bone marrow metastasis from the same patient before and after treatment, respectively, and are both highly susceptible to ZIKV. Thus, Table 1 identifies that ZIKV can target neuroblastoma cells which originate from the primary tumour, metastatic sites and metastatic sites which are resistant to standard neuroblastoma therapy. SK-N-AS, T-268 and JFEN are highly resistant (cell viability >80%) to ZIKV infection. Susceptibility is independent of patient sex, cell line origin, morphology and MYC-N status (Table 1). The non-sympathetic nervous system and non-paediatric origin of the T-268 and JFEN cells likely explain their resistance to ZIKV infection, as ZIKV has a tropism for paediatric nervous system cancer cells. The resistance of the paediatric SK-N-AS cell line is governed by CD24 expression, which regulates the basal antiviral state of these cells (13, 19). In conclusion, our analysis demonstrates that ZIKV is a strong candidate to employ against paediatric neuroblastoma as oncolytic virotherapy.

**Table 1.** ZIKV infects a multitude of neuroblastoma cell lines. Cell lines are ranked by the degree to which ZIKV infection reduces their cell viability. Ranking was determined by taking the percentage to which ZIKV either reduced cell viability or induced cell death (depending on the in vitro assay used in the study) and applying a ranking system from 1 to 5, with higher numbers denoting a strong ability of ZIKV to reduce cell viability.

| Cell line | Cell Viability | Age (years) | Sex | Cancer Type | Cell line origin | Morphology | MYCN Status | References |
| --- | --- | --- | --- | --- | --- | --- | --- | --- |
| SH-SY5Y | 5 | 4 | Female | Neuroblastoma | Bone marrow metastasis (thorax) | Epithelial | non-amplified | (50,51) |
| SK-N-SH | 5 | 4 | Female | Neuroblastoma | Bone marrow metastasis (thorax) | Epithelial | non-amplified | (21,52) |
| SK-N-BE(2) | 5 | 2 | Male | Neuroblastoma | Bone marrow metastasis | Neuroblast | amplified | (50,51) |
| SK-N-BE(2)-M17 | 5 | 2 | Male | Neuroblastoma | Bone marrow metastasis | Neuroblast | amplified | (51) |
| SK-N-DZ | 5 | 2 | Female | Neuroblastoma | Bone marrow metastasis | Epithelial | amplified | (50,51) |
| IMR-32 | 5 | 1 | Male | Neuroblastoma | Abdominal mass primary tumour | Neuroblast, Fibroblast | amplified | (13,50,51) |
| SMS-KAN | 5 | 3 | Female | Neuroblastoma | Pelvic primary tumour | Neuroblast | amplified | (13) |
| SMS-KCNR | 5 | 1 | Male | Neuroblastoma | Bone marrow metastasis (adrenal) | Neuroblast | amplified | (50) |
| SK-N-FI | 5 | 11 | Male | Neuroblastoma | Bone marrow metastasis | Epithelial | non-amplified | (50) |
| CHLA-42 | 5 | 1 | NA | Neuroblastoma | Bone marrow metastasis | Epithelial | non-amplified | (13) |
| SK-N-BE(1) | 4 | 2 | Male | Neuroblastoma | Bone marrow metastasis | Neuroblast | amplified | (13) |
| LA-N-6 | 3 | 5 | Male | Neuroblastoma | Bone marrow metastasis (adrenal) | Neuroblast | non-amplified | (13) |
| SK-N-AS | 1 | 6 | Female | Neuroblastoma | Bone marrow metastasis (adrenal) | Epithelial | non-amplified | (13) |
| T-268 | 1 | 22 | Female | Olfactory neuroblastoma | Metastasis (paraspinal mass) | NA | NA | (50) |
| JFEN | 1 | 22 | Male | Olfactory neuroblastoma | Metastasis (chest wall) | NA | NA | (50) |

| Ranking System | Cell Viability |
| --- | --- |
| 5 | <20% |
| 4 | 20-40% |
| 3 | 40-60% |
| 2 | 60-80% |
| 1 | >80% |

### ZIKV strains possess different therapeutic potential against neuroblastoma cells

The neurotropism of the Asian lineage makes it the clear choice, over the African lineage, for developing ZIKV oncolytic virotherapy against brain tumours. However, the situation for Neuroblastoma is not so apparent. Independent studies have demonstrated inherent differences in the ability of varying ZIKV strains to infect, replicate, and kill neuroblastoma cells (20, 21). Here, we assess all published data concerning ZIKV infection of neuroblastoma cells and ranked the viral strains based on their ability to infect neuroblastoma cells, produce fresh viral progeny and reduce cell viability (Table 2). We identify the PRVABC59 Asian, Uganda #976 African and MR766 African strains as the top three candidates (Table 2). Interestingly, the PRVABC59 Asian strain induces significantly more DEGs and splice events in SH-SY5Y cells compared to the African MR766 strain; including more immune and inflammatory response genes (22). Brain metastases develop in 5-11% of neuroblastoma patients and are correlated with poor prognosis (23). The neurotropism of the Asian lineage may enhance the therapeutic potential of ZIKV by targeting these brain metastases. The multiple ZIKV pandemics identified that infection by an Asian strain is generally well accommodated by children, thus providing evidence for the safety of employing an Asian strain. Consequently, from those tested to date, we identify that the PRVABC59 strain demonstrates the greatest promise for development as oncolytic virotherapy against paediatric neuroblastoma.

**Table 2.**
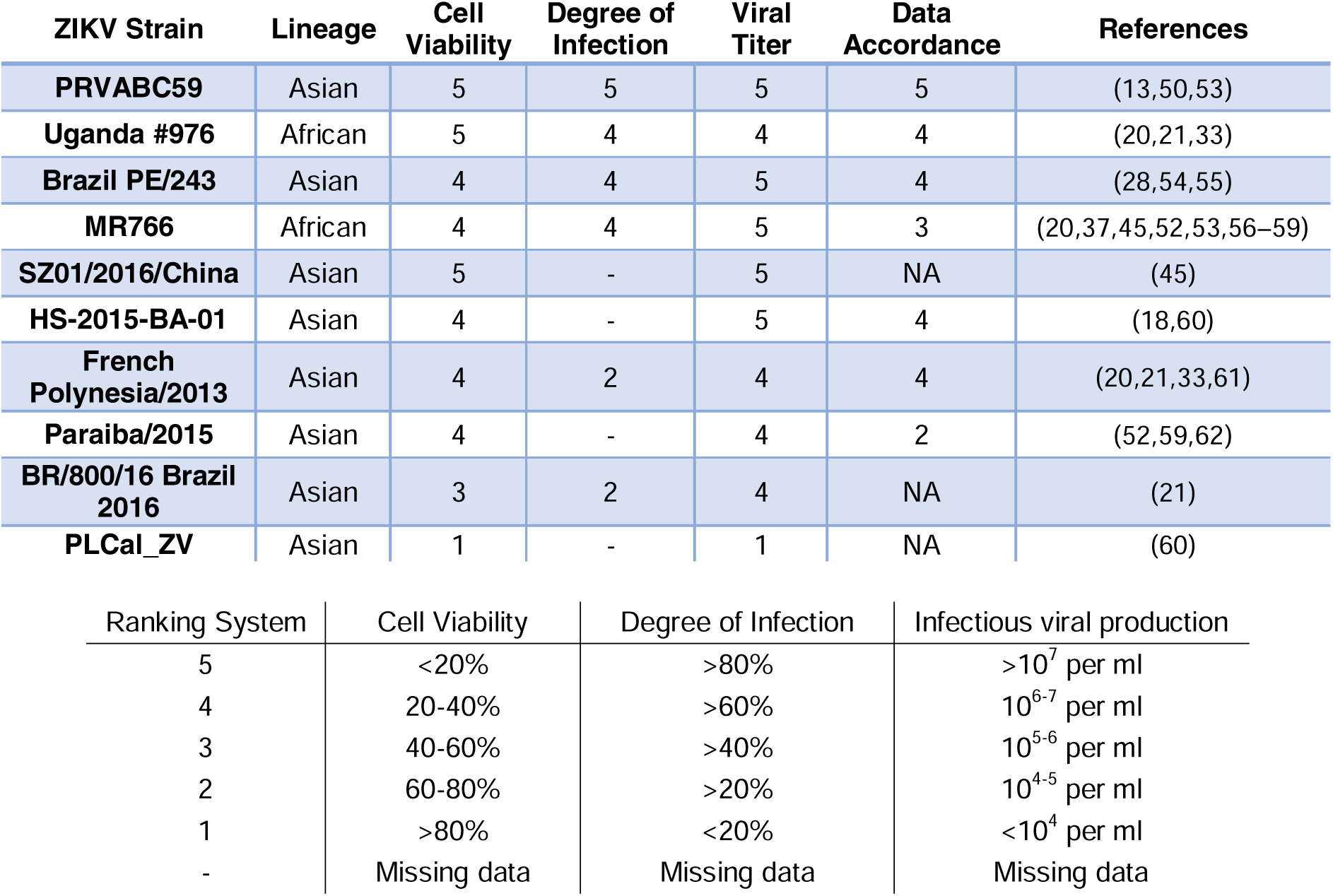
ZIKV strains possess differing therapeutic potential against neuroblastoma cells. ZIKV strains are ranked by their ability to infect, replicate within and significantly reduce the cell viability of a multitude of neuroblastoma cells. Data accordance denotes the degree of similarity of the results between publications which performed cell viability of cell death assays in neuroblastoma cells using the same ZIKV strain. Data accordance of 5 denotes that the findings of one publication closely support the findings from another, a data accordance of 1 denotes publications reporting vastly contrasting results. When a viral strain is published in only one paper, it is allocated a data accordance of NA.

### ZIKV infection of neuroblastoma cells induces changes at the transcriptome level

Differential gene expression analysis identifies 453 and 256 significantly upregulated and downregulated genes (fold change > 1.5), respectively, in ZIKV-infected paediatric neuroblastoma SH-SY5Y cells (Figure 1A). Gene Ontology (GO), Reactome and KEGG pathway analysis identifies nine significantly upregulated and 12 significantly downregulated terms (Figure 1B-C). Upregulated processes include “TNF signalling”, lipid metabolism (“Cholesterol biosynthesis”, “Cholesterol biosynthetic process”, “Activation of gene expression by SREBF (SREBP)”), ER stress (“Response to endoplasmic reticulum stress”, “XBP1(S) activates chaperone genes”) and transcription (“BMAL1:CLOCK, NPAS2 activates circadian gene expression”, “Positive regulation of transcription from RNA polymerase II promoter”). The downregulated terms are predominantly cell cycle- and DNA replication-related processes and this downregulation is apparent when the “Cell Cycle” KEGG pathway is plotted for all DEGs (fold change > 0) (Figure 1D). A potential explanation for this observation is that ZIKV can disrupt the cell cycle by targeting the centrioles in neuroblastoma cells (24).

**Figure 1.**
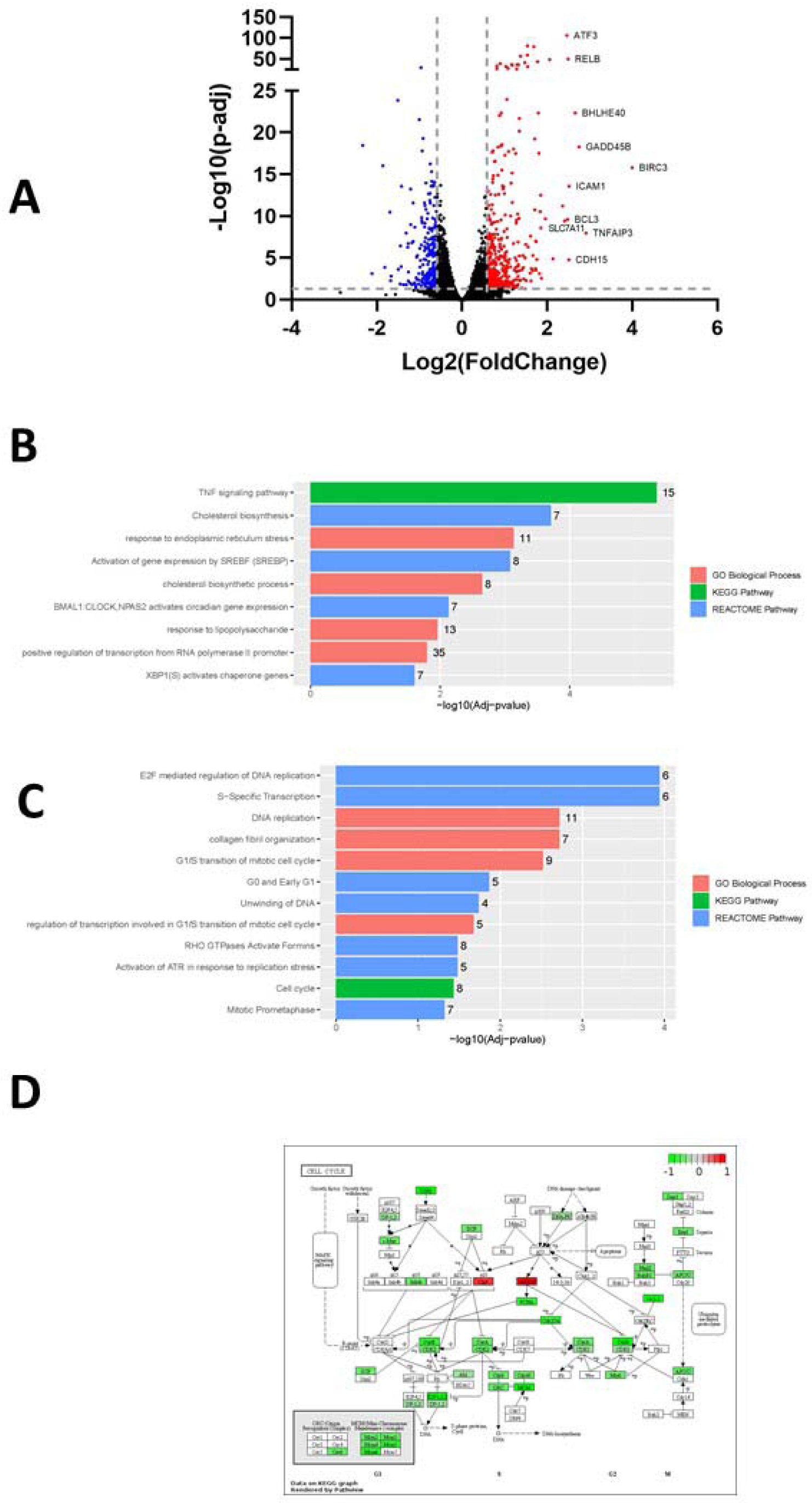
Differential gene expression, Gene Ontology (GO) and Pathway Analysis of ZIKV infection in SH-SY5Y cells. Volcano plot of genes differentially expressed in response to ZIKV infection of SH-SY5Y cells, with the top ten upregulated genes labelled (A). Significantly up-regulated (B) and down-regulated (C) GO Biological Processes, KEGG and Reactome pathways in response to ZIKV infection in SH-SY5Y neuroblastoma cells. KEGG map of the Cell Cycle, plotted using all DEGs (fold change > 0) (D). Significance values are corrected for multiple testing using the Benjamini and Hochberg method (padj < 0.05).

### ZIKV induces TNF Signalling in neuroblastoma cells

Of the top ten upregulated DEGs in SH-SY5Y cells, four (BIRC3, TNFAIP3, ICAM1 and BCL3) are components of the TNF signalling pathway (Figure 1A). The TNF pathway is particularly interesting to consider for oncolytic virotherapy since it may play a role in both oncolysis (direct cell death) and the anti-tumoral immune response. Here, mapping the “TNF Signalling” KEGG pathway for ZIKV-infected SH-SY5Y cannot deduce if ZIKV may activate CASP-mediated apoptosis or CASP-independent necroptosis (Figure 2A). However, ZIKV-infected SH-SY5Y cells clearly show significant upregulation of transcription factors (AP-1, cEBPβ and CREB), leukocyte recruitment and activation (CCL2 and CSF1), intracellular signalling (BCL3, NFKBIA, TNFAIP3 and TRAF1) and cell adhesion genes (Icam1 and Vcam1) (Figure 2A). ZIKV significantly upregulates the expression of multiple Activator protein 1 (AP-1) transcription factors, including members from all four AP-1 subfamilies (ATF, JUN, FOS and MAF) in SH-SY5Y cells (Figure 2B). AP-1 can regulate the expression of a diverse set of genes in response to nutrients, cytokines, stress or pathogen infection, and is involved in innate and adaptive immunity, differentiation, proliferation, survival and apoptosis (25). AP-1 increases the range of genes that it can regulate through interaction with additional factors, including NF-κB. NFKB1 and NFKB2 are both significantly upregulated by ZIKV infection (data not shown). AP-1 transcription factors can regulate the immune response of tumours, and significant AP-1 upregulation by ZIKV infection potentially identifies AP-1 as a mechanism through which ZIKV could yield an anti-tumoral immune response against neuroblastoma *in vivo* (26).

**Figure 2.**
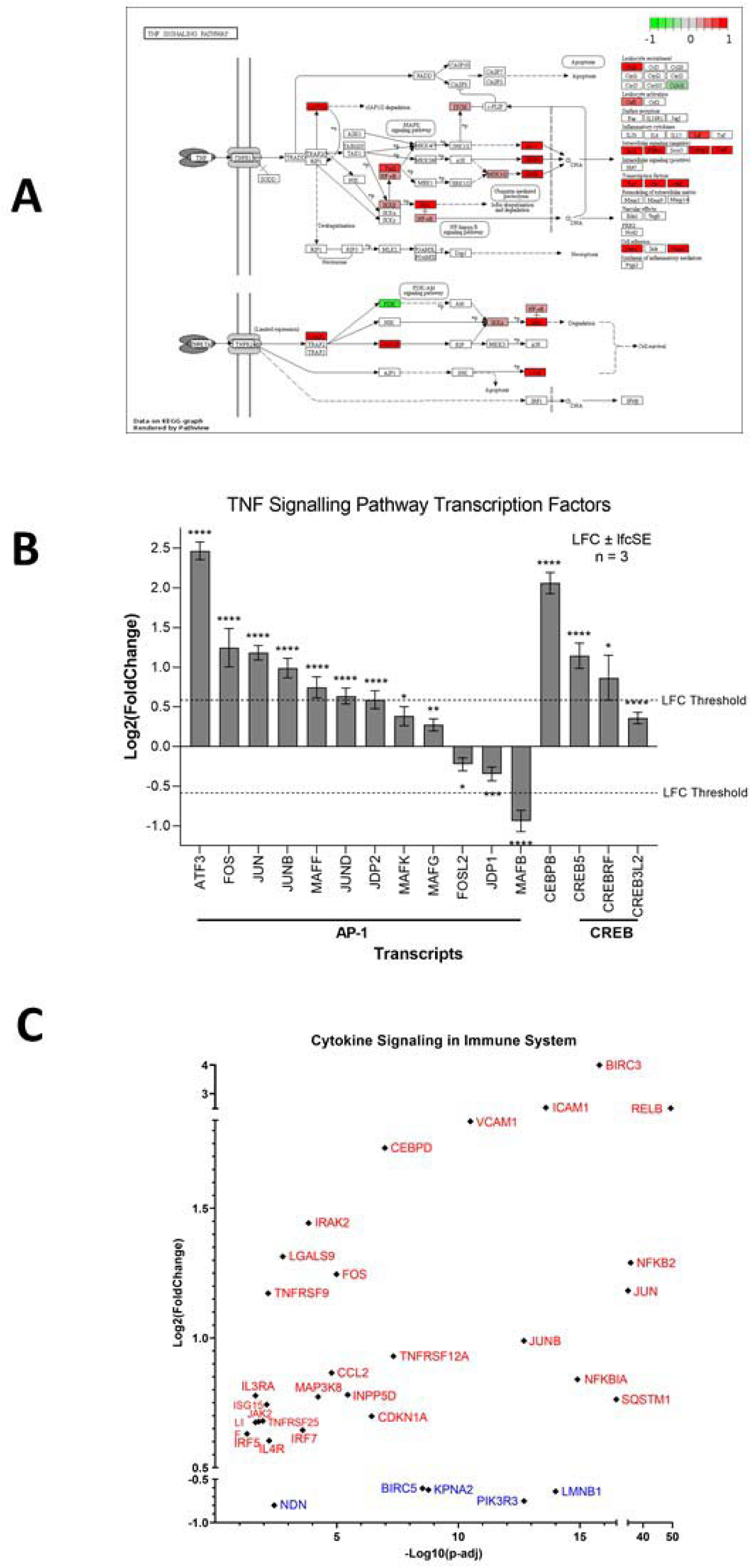
ZIKV infection upregulates the TNF Signalling pathway in neuroblastoma cells. KEGG Map plotted for the TNF Signalling Pathway, plotted using all DEGs (fold change > 0) (A). Expression levels of TNF Signalling Transcription Factors in SH-SY5Y cells in response to ZIKV infection (B). Expression levels of Cytokine Signalling in Immune System genes in SH-SY5Y cells in response to ZIKV infection (C). Significance values are corrected for multiple testing using the Benjamini and Hochberg method (padj < 0.05). A threshold line of Log_2_(1.5 Fold Change) has been applied for the expression values. Log_2_FoldChange (LFC) ± standard error of the LFC estimate (lfcSE), n = 3.

Here, we identify CCL2 (MCP-1) to be significantly upregulated by ZIKV infection, and two independent studies have shown CCL2 to be secreted by ZIKV-infected SH-SY5Y cells (27, 28). CCL2 is a pro-inflammatory mediator that recruits leukocytes via chemotaxis to infiltrate tissues, including the CNS, to stimulate inflammation (29). A non-neurotoxic HSV-based oncolytic virotherapy, engineered to express physiologically relevant levels of CCL2 (M010), significantly reduced Neuro-2a neuroblastoma growth in the flank of immune-competent mice and recruited CD4+ and CD8+ T-cells to infiltrate the tumour (29). Additionally, CCL2 is secreted by ZIKV-infected cultured canine glioblastoma cells when in the presence of monocytes and is detected in serum and CSF samples of canines bearing spontaneous brain tumours following ZIKV infection (12). We propose here that CCL2 may be capable of inducing an anti-tumoral immune response against paediatric neuroblastoma during ZIKV infection. Supporting the notion of a ZIKV-induced inflammatory response, 32 genes implicated in cytokine signalling in the immune system are significantly differentially expressed in ZIKV-infected SH-SY5Y cells: 27 upregulated and 5 downregulated (Figure 2C).

### ZIKV induces lipid metabolism in neuroblastoma cells

ZIKV infection significantly upregulates lipid metabolism-related terms in SH-SY5Y cells; specifically “Cholesterol biosynthesis” and “Activation of gene expression by SREBF (SREBP)” (Figure 1B). Cholesterol and lipids are essential cellular components and there are complex systems that function to regulate their intracellular abundance and localisation. These systems include regulation of cholesterol biosynthesis by the SREBP pathway, intracellular cholesterol trafficking, and cholesterol efflux by the LXR pathway. Cholesterol and fatty acids are required for multiple stages of the flaviviral life cycle, including regulating viral entry, the formation of viral replication complexes in the ER membrane and viral egress (30). ZIKV elevates lipogenesis and remodels the composition of the lipid classes in infected SK-N-SH cells (31). Here, we identify several approaches to regulate ZIKV infection of neuroblastoma cells through modification of intracellular lipid levels (Table 3). These include supplementation with pathway regulators (PF-429242, Fenofibrate, Lovastatin, U18666A and LXR 623) or exogenous lipids (Oleic acid, DHA and Cholesterol).

**Table 3.** Compounds capable of regulating ZIKV infection in neuroblastoma cells through modifying lipid abundance, composition and localisation.

| Compound | Cell Line | Mechanism of Action | Effect on ZIKV infection | References |
| --- | --- | --- | --- | --- |
| Bafilomycin A1 (V-ATPase inhibitor) | SH-SY5Y | Impairs acidification of endosomal-lysosomal compartments | Restrict | (33) |
| U18666A | SH-SY5Y | Cholesterol accumulation impairs late endosomes & lysosomes | Restrict | (33) |
| LXR 623 (LXR pathway agonist) | SK-N-SH | Induces cholesterol efflux | Restrict | (34) |
| PF-429242 (SREBP pathway inhibitor) | SK-N-SH | Reduces intracellular lipid levels | Restrict | (31) |
| Fenofibrate (SREBP pathway inhibitor) | SK-N-SH | Reduces intracellular lipid levels | Restrict | (31) |
| Lovastatin (SREBP pathway inhibitor) | SK-N-SH | Reduces intracellular lipid levels | Restrict | (31) |
| Oleic Acid | SK-N-SH | Increases lipid droplet abundance | Enhance | (31) |
| Cholesterol | SK-N-SH | Increases lipid droplet abundance | Restrict | (31) |
| DHA | SH-SY5Y | Anti-inflammatory and neuroprotective effects against ZIKV infection | Restrict | (28) |

The SREBP pathway is a principal regulator of fatty acid and cholesterol biosynthesis. The SREBF1 and SREBF2 transcription factors control this pathway, and although they share a small degree of redundancy, they primarily regulate the expression of fatty acid biosynthesis and cholesterol biosynthesis target genes, respectively (32). Both SREBF2 and SREBF2-AS1 are significantly upregulated in ZIKV-infected SH-SY5Y cells (Figure 3A) and pathway analysis identifies significant upregulation of “Cholesterol biosynthesis”. Three SREBP pathway inhibitors (PF-429242, Fenofibrate and Lovastatin) reduce the capability of ZIKV to infect SK-N-SH neuroblastoma cells (Table 3). We identify here that the SREBP pathway is essential for ZIKV infection and propose that it may upregulate cholesterol biosynthesis through the SREBF2 transcriptional pathway in neuroblastoma cells.

**Figure 3.**
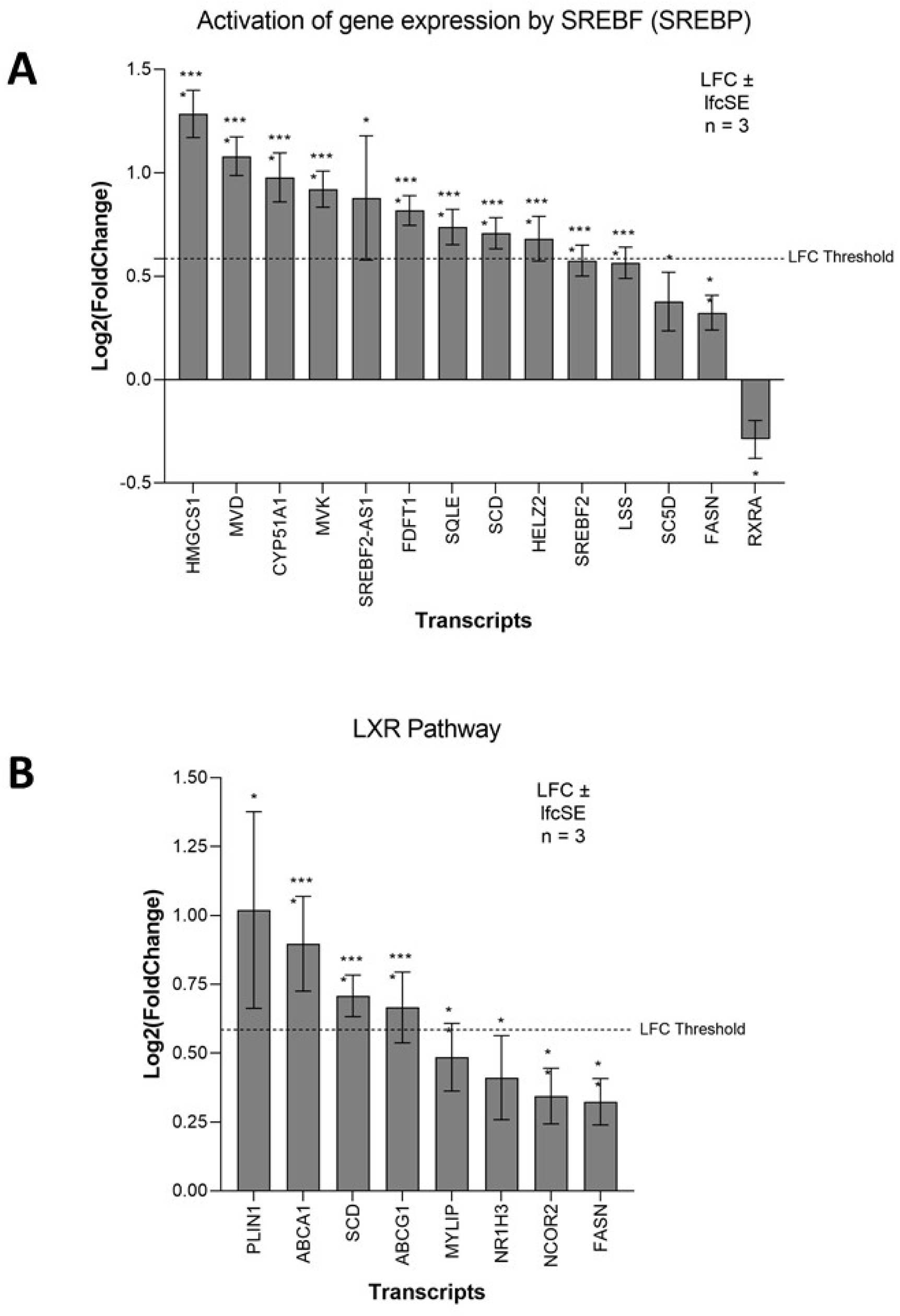
ZIKV infection upregulates lipid metabolism in neuroblastoma cells. Expression of SREBP pathway (A) and LXR pathway (B) genes in neuroblastoma cells in response to ZIKV infection. Significance values are corrected for multiple testing using the Benjamini and Hochberg method (padj < 0.05). A threshold line of Log_2_(1.5 Fold Change) has been applied for the expression values. Log_2_FoldChange (LFC) ± standard error of the LFC estimate (lfcSE), n = 3.

Both U18666A and exogenous cholesterol restricts ZIKV infection of neuroblastoma cells (Table 3). U18666A is an intracellular cholesterol transport inhibitor that causes the accumulation of cholesterol in lysosomes to hinder the endosomal-lysosomal system (33). Exogenous cholesterol also leads to the inactivation of the late endosomal-lysosomal compartments through a build-up of cholesterol (33). Collectively, this identifies a dependence of the ZIKV life cycle in neuroblastoma cells on intracellular cholesterol for the correct functioning of the endosomal-lysosomal system.

The Liver X receptor (LXR) pathway agonist (LXR 623) promotes cholesterol efflux (Table 3). Flaviviruses require cholesterol for the restructuring of host membranes, and LXR 623 demonstrates this dependence of ZIKV in neuroblastoma by preventing ZIKV-induced vesicle production and ER expansion in SK-N-SH cells (34). The LXR pathway and expression of its downstream lipid homeostasis genes are regulated by the transcription factors LXR-α (NR1H3) and LXR-β (NR1H2). LXR-α protein is significantly increased by ZIKV infection of SK-N-SH neuroblastoma cells from 48hr (34). Although LXR-α mRNA is only marginally upregulated in our study, two major cholesterol efflux factors that are downstream of the LXR pathway, ATP-Binding Cassette A1 and G1 (ABCA1 and ABCG1), are significantly upregulated (Figure 3B). This identifies a dependence of ZIKV on the LXR pathway and suggests that ZIKV may manipulate this pathway in neuroblastoma cells to upregulate cholesterol efflux.

We identify here that lipid abundance, localisation, trafficking and metabolism regulate ZIKV infection of neuroblastoma cells and that ZIKV likely remodels the cellular lipid composition to help produce a favourable environment for efficient replication.

### ZIKV induces and is dependent on the ER stress response in neuroblastoma cells

ZIKV upregulates ER-stress-related terms in SH-SY5Y cells, principally “Response to endoplasmic reticulum stress” and “XBP1(S) activates chaperone genes” (Figure 1B). The Unfolded Protein Response (UPR) dictates the ER-stress response. The UPR is normally inactive due to the ER chaperone BIP sequestering three ER stress sensors (IRE1, PERK and ATF6). Under stress conditions BIP releases IRE1, PERK and ATF6 to assist protein folding, allowing them to activate their respective UPR-mediated ER-stress pathways.

Activation of the IRE1-mediated UPR leads to IRE1 splicing a 26bp region from the ubiquitously expressed XBP1 mRNA. The active transcription factor XBP1(S) then drives the expression of genes to help alleviate ER stress, primarily chaperone and ER-associated protein degradation (ERAD) genes. ZIKV infection significantly upregulates 15 genes of the IRE1-mediated “XBP1(S) activates chaperone genes” Reactome pathway in SH-SY5Y cells (Figure 4A). XBP1 is the most highly upregulated gene and others include the ERAD gene SYVN1 and the chaperones DNAJC3 and DNAJB9 (Figure 4A). Multiple IRE1-mediated UPR genes (EDEM1, SYVN1, SSR1, SRPRB, ATP6V0D1, and EXTL3) are ZIKV dependency factors in hiPSC-NPCs (Supplementary data). Chemical inhibition of IRE1 by 4μ8C impairs ZIKV infection *in vivo* (35). Here, we identify that ZIKV significantly upregulates the IRE1-mediated UPR in SH-SY5Y cells and that ZIKV is dependent on this for efficient infection, likely as a means to regulate and combat viral replication-induced ER stress.

**Figure 4.**
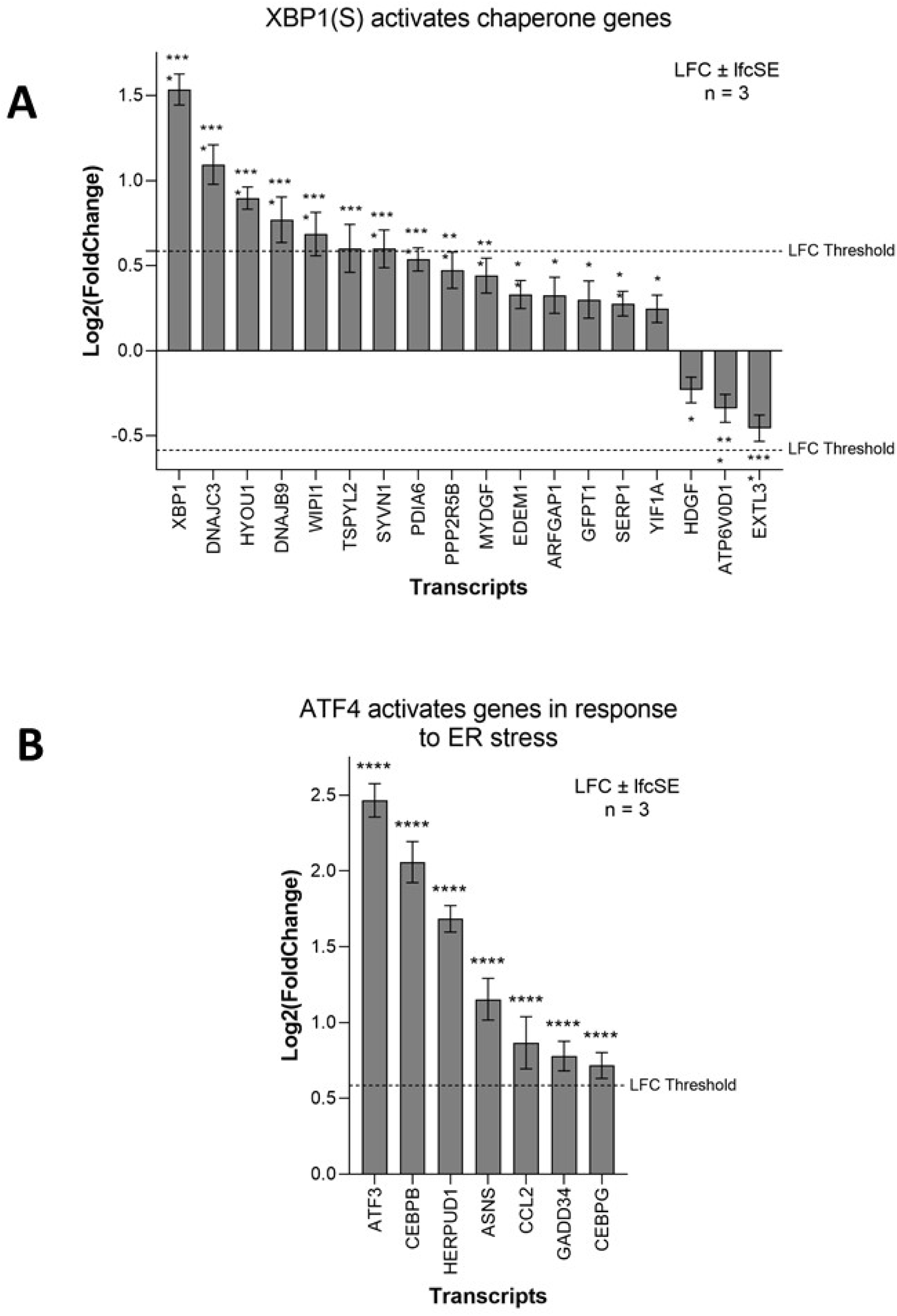
ZIKV infection activates the UPR in neuroblastoma cells. Expression levels of the XBP1(S) activates chaperone genes (A) and ATF4 activates genes in response to endoplasmic reticulum stress (B) genes in neuroblastoma cells in response to ZIKV infection. Significance values are corrected for multiple testing using the Benjamini and Hochberg method (padj < 0.05). A threshold line of Log_2_(1.5 Fold Change) has been applied for the expression values. Log_2_FoldChange (LFC) ± standard error of the LFC estimate (lfcSE), n = 3.

PERK-mediated UPR regulates the expression of genes involved in apoptosis, redox, amino acid transport and autophagy through eIF2 phosphorylation and the transcription factor ATF4. ZIKV infection significantly upregulates seven genes of the Reactome pathway “ATF4 activates genes in response to endoplasmic reticulum stress” (Figure 4B), including the ERAD gene HERPUD1 and the transcription factors ATF3, CEBPB and CEBPG. GADD34 (PPP1R15A) which usually dephosphorylates eIF2α in a negative feedback loop, is significantly upregulated here by ZIKV infection, and ZIKV induces eIF2 phosphorylation in SK-N-SH cells (36). ZIKV likely upregulates GADD34 to combat ER stress-induced translational repression, as fresh virions require de novo protein synthesis. CHOP (DDIT3) is a pro-apoptotic protein downstream of the PERK UPR pathway that others have observed to be significantly upregulated in SH-SY5Y and SK-N-SH cells in response to ZIKV infection (36–38). CHOP induces apoptotic markers, including Caspase 3, leading to cell death. Interestingly, multiple PERK-mediated UPR genes (ATF4, EIF2AK1, EIF2AK2, EIF2AK3 and EIF2AK4) are ZIKV dependency factors in hiPSC-NPCs (Supplementary data).

We conclude that ZIKV specifically upregulates and is dependent on the IRE1 and PERK branches, but not the ATF6 branch, of the UPR ER stress response in SH-SY5Y cells; conclusions that are supported by others (36, 37).

### ZIKV is dependent on the EMC in neuroblastoma cells

To determine what host mechanisms ZIKV may be dependent on, we cross-referenced the 22 known proteins which ZIKV requires to infect neuroblastoma cells with ZIKV dependency factors from four cell lines (Figure 5A, Supplementary data). Between 72-94% of the dependency factors identified across the five different cell lines are cell-specific, highlighting how ZIKV utilises differing host factors across different cell types for its life cycle. The sparse overlap of ZIKV dependency factors identifies only one factor common to all five cell types. MMGT1 (EMC5) is a key component of the Endoplasmic Reticulum (ER) Membrane Protein Complex (EMC). The EMC is a hetero-oligomer composed of ten subunits, has chaperone properties by assisting multi-transmembrane protein folding, and is implicated in ER stress, flavivirus infection and lipid trafficking (39). To assess if ZIKV is dependent on additional EMC subunits during infection, we searched for them in our ZIKV dependency factor dataset. We identify that ZIKV has a strong dependence on the EMC independent of cell type; all ten EMC proteins are ZIKV-dependency factors in hiPSC-NPC, eight are in HeLa and three in GSC and HEK293FT cells (Figure 5B). The EMC facilitates the expression of ZIKV transmembrane proteins (NS2B, NS4A and NS4B), ZIKV NS4B interacts with EMC subunits, and disrupting the EMC impedes infection by ZIKV and other flaviviruses (40, 41). We propose that the EMC stabilises ZIKV proteins through integration into the ER membrane, thus permitting efficient infection in neuroblastoma cells. If investigated, we predict that additional EMC subunits would present as ZIKV dependency factors in neuroblastoma cells.

**Figure 5.**
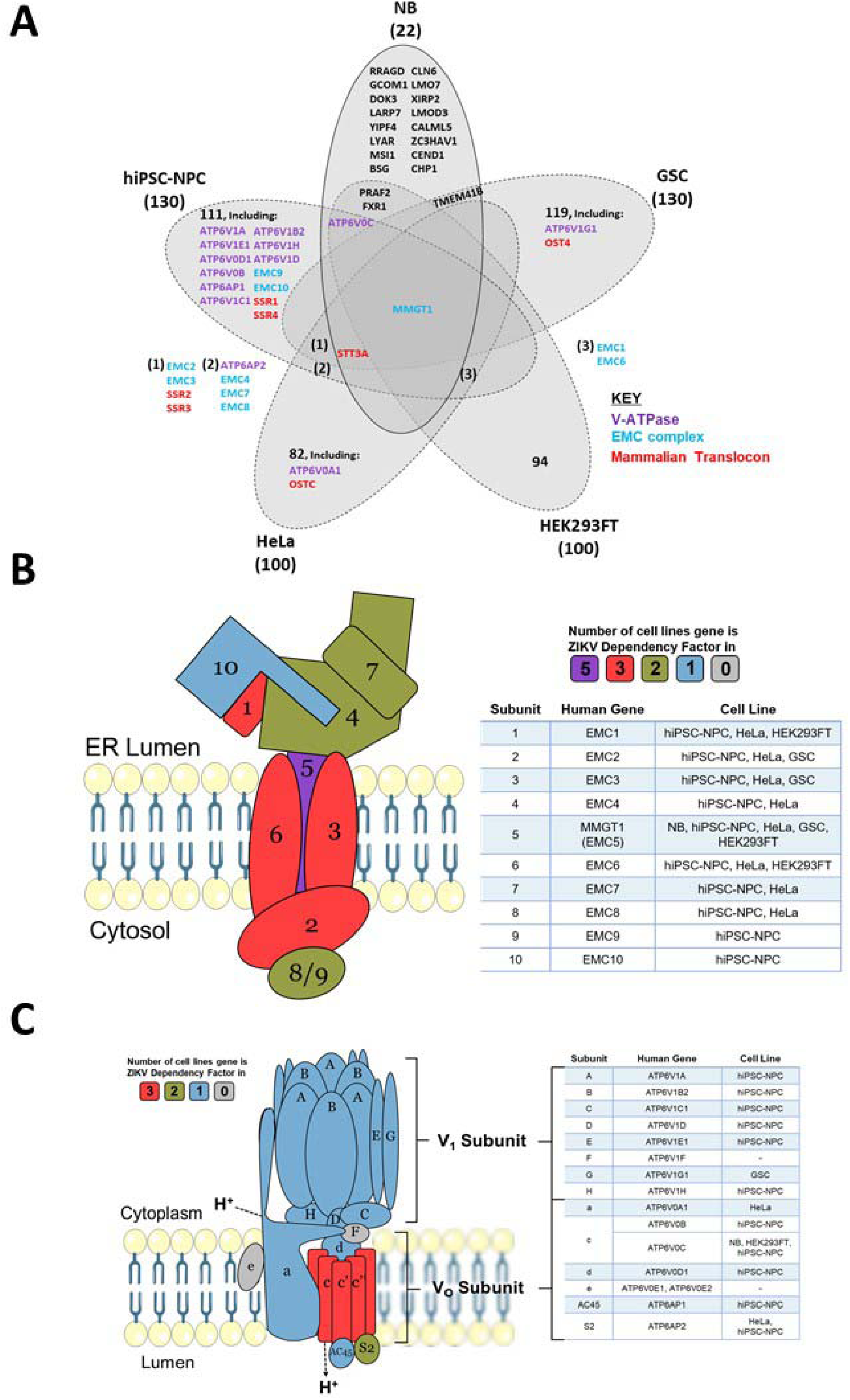
ZIKV dependency factors. Venn diagram of known ZIKV dependency factors across Neuroblastoma (NB), Glioma Stem Cells (GSC), hiPSC-NPC (NPC), HEK293FT and HeLa cells, to identify shared and cell-specific factors and protein complexes which ZIKV is dependent on for infection (A). Diagram of the Endoplasmic Reticulum (ER) Membrane Protein Complex (EMC) (B) and V-ATPase (C), based on their crystal structures. For the subunits in B and C, colours are allocated based on the number of cell types in which they act as ZIKV dependency factors (cell types stated in the adjacent tables).

### ZIKV is dependent on the V-ATPase in neuroblastoma cells

Acidification of the endosomal-lysosomal system by the V-ATPase is a property which viruses can utilise to drive the release of their nucleocapsid into the cytosol. ATP6V0C, a central component of the V-ATPase, is a ZIKV-dependency factor in neuroblastoma, hiPSC-NPC and HEK293FT cells (Figure 5A). Twelve additional V-ATPase subunits are ZIKV dependency factors across GSC, hiPSC-NPC and HeLa cells (Figure 5C). These 13 genes consist of multiple subunits from the Vo proton translocation and V1 ATP hydrolytic domains, identifying a functional dependence of ZIKV on the entire V-ATPase complex. The V-ATPase inhibitor Bafilomycin A1 specifically binds ATP6V0C and through V-ATPase inhibition prevents lysosomal acidification and the autophagy-lysosome pathway (42). Bafilomycin A1 inhibits ZIKV infection of SH-SY5Y cells, supporting our observation (Table 3). siRNA silencing of the V-ATPase significantly impairs ZIKV infection of T98G glioblastoma cells, collaborating its requirement for infection of nervous system tumour cells (43). We propose that loss of V-ATPase function impairs ZIKV infection in SH-SY5Y cells due to a perturbed pH gradient in the endosomal system. This likely prevents fusion of the viral envelope with the endosomal membrane for release of the nucleocapsid, therefore, trapping ZIKV for degradation in the lysosome, as observed in Vero cells (44).

### ZIKV NS4B possesses oncolytic capability against neuroblastoma cells

ZIKV NS4B protein is principally responsible for the oncolytic effect in SH-SY5Y cells, via activating the mitochondrial apoptotic pathway (45). Determining the interactions and mechanisms underpinning this may yield opportunities to develop a paediatric neuroblastoma therapy based on ZIKV NS4B. ZIKV NS4B has 130 known host interaction partners in SK-N-BE(2) neuroblastoma cells (14). Here, we analyse this interactome and identify multiple pathways which we, and others, have implicated during ZIKV infection of neuroblastoma cells: including mitochondrial-, lipid metabolism- and ER-associated processes (Figure 6). ZIKV NS4B interacts with ten lipid biosynthesis proteins, three (SCD, SC5D and DHCR7) of which are expressed in response to SREBP pathway activation. This identifies a direct interaction between ZIKV and host lipid metabolism and the SREBP pathway, supporting our previous observations at the transcriptome level.

**Figure 6.**
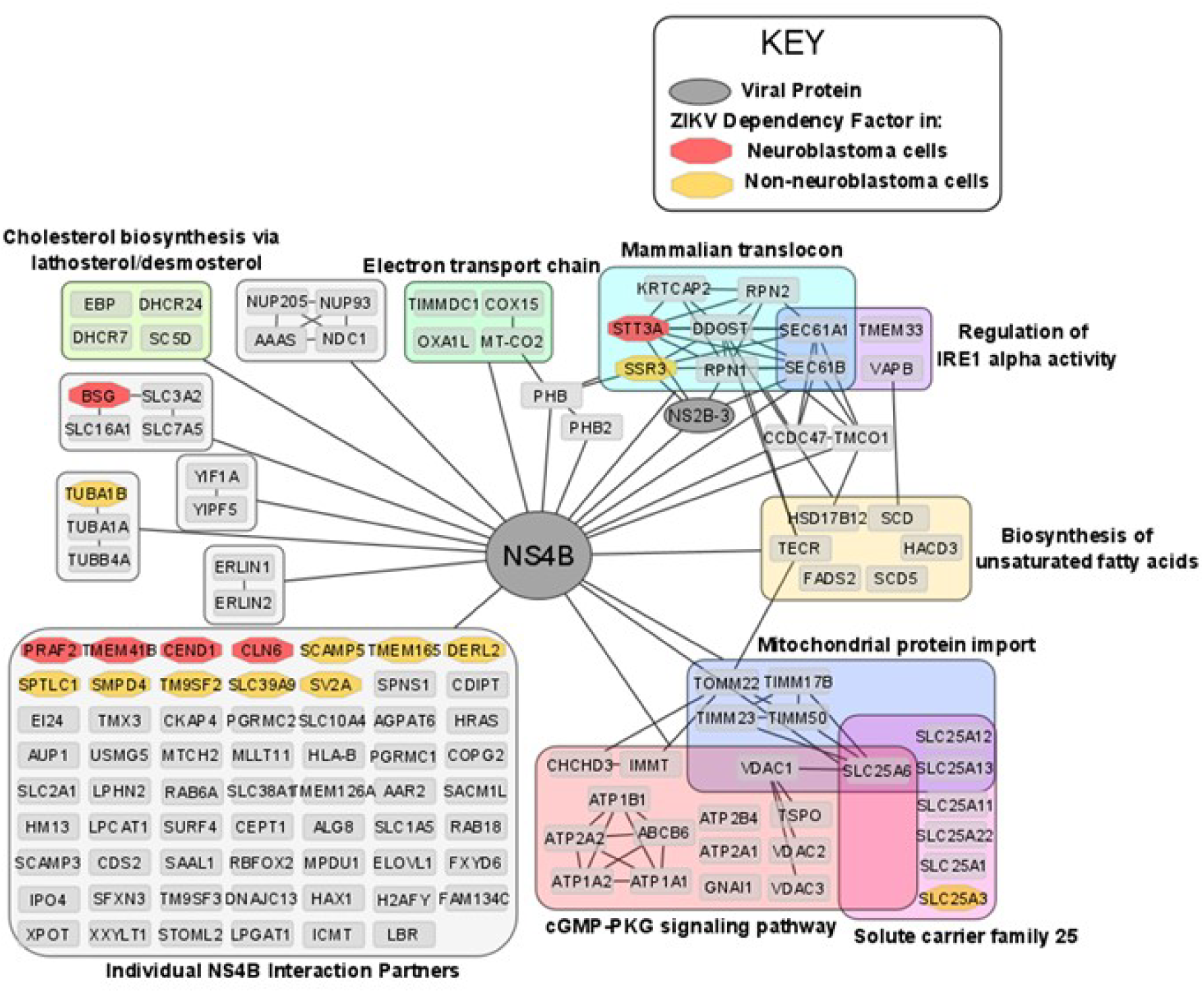
The ZIKV NS4B interactome in neuroblastoma cells and its interaction with ZIKV dependency factors. Nodes are grouped and labelled according to any sets of high-confidence interactions between the host proteins, and by pathways they are involved in. Cross-referencing the interactome with the ZIKV dependency factor datasets identifies the interaction of NS4B with dependency factors in neuroblastoma cells (red) and in GSC, hiPSC-NPC, HeLa and HEK293FT cells (collectively termed non-neuroblastoma cells, orange). To aid visualisation, any nodes possessing no high confidence interactions, other than their interaction with NS4B, have been grouped. All nodes within Figure 6 interact with NS4B, thus, to aid visualisation all edges between NS4B and nodes within a group have been condensed into a single edge between NS4B and the grouped set of nodes.

ZIKV NS4B recruits BAX to the mitochondria, triggers its activation, and releases Cytochrome c from mitochondria to induce mitochondrial cell death in SH-SY5Y cells (45). ZIKV NS4B interacts with a multitude of mitochondrial genes (Figure 6). Including, electron transport chain proteins (TIMMDC1, MT-CO2, COX15 and OXA1L), mitochondrial translocases that import proteins into the mitochondrial matrix (TOMM22, TIMM23, TIMM50 and TIMM17B) and Solute Carrier Family 25 members for transport of solutes across the mitochondrial membrane (SLC25A1, SLC25A3, SLC25A6, SLC25A11, SLC25A12, SLC25A13 and SLC25A22). Specifically, MT-CO2, COX15 and OXA1L are conserved catalytic core, assembly and accessory subunits of the Cytochrome c oxidase complex, respectively. The Cytochrome c oxidase complex tightly couples Cytochrome c to the inner mitochondrial membrane. We propose that NS4B interacts with Cytochrome c oxidase to uncouple it from Cytochrome c, causing Cytochrome c release through the BAX pore into the cytosol to drive the mitochondrial cell death pathway in neuroblastoma cells.

### ZIKV NS4B interacts with the Mammalian Translocon in neuroblastoma cells

ZIKV NS4B interacts with and is dependent on multiple proteins of the Mammalian Translocon (Figures 5A & 6). The mammalian translocon is primarily composed of the Oligosaccharyl Transferase (OST) complex, the Sec61 complex and the translocon-associated protein (TRAP) complex (46). The multimeric OST complex co-translationally N-glycosylates proteins within the ER to assist protein folding, stability and trafficking. The Sec61 complex, a heterotrimer of Sec61α, Sec61β and Sec61γ, co-translationally translocates newly synthesised proteins across the ER and during ER stress can regulate IRE1α activity. TRAP is a heterotetramer of SSR1, SSR2, SSR3 and SSR4 that assists co-translational translocation of proteins into the ER and can prevent aberrant N-linked glycosylation during ER stress.

STT3A, a principal component of the OST complex, interacts with ZIKV NS4B and is a ZIKV dependency factor in neuroblastoma, GSC, hiPSC-NPC and HeLa cells (Figures 5A & 6). Regarding the additional OST subunits, ZIKV NS4B interacts with DDOST, RPN1, RPN2 and KRTCAP2, and OSTC and OST4 are ZIKV dependency factors in HeLa cells and GSCs, respectively (Figures 5A & 6). Two forms of the OST complex exist, the STT3A and STT3B OST paralogs. KRTCAP2 and OSTC are STT3A-specific OST factors, which permit interaction of STT3A with the translocon, whilst TUSC3 and MAGT1 are STT3B-specific OST factors (46). STT3A and both STT3A-specific OST factors are ZIKV interaction partners and/or dependency factors, but neither STT3B nor the STT3B-specific OST factors are. In addition to the OST factors, multiple N-linked glycosylation-related proteins (DPM1, DERL3, SYVN1, UBE2G2 and UBE2J1) are also ZIKV dependency factors in non-neuroblastoma cells (Supplementary data). The OST complex inhibitor NGI-1 blocks ZIKV infection of Huh7 cells, and disrupting ZIKV prM and E protein N-glycosylation impairs the release of infectious ZIKV particles from Vero cells (47, 48). Here, we identify that STT3A functions as a bonafide ZIKV dependency factor in multiple cell types, and conclude that efficient infection of neuroblastoma cells by ZIKV is likely dependent on the STT3A OST paralog for N-glycosylation of its viral proteins.

ZIKV NS4B interacts with SEC61A1 and SEC61B of the Sec61 complex in neuroblastoma cells and the Sec61α inhibitor Mycolactone impedes ZIKV infection of HeLa cells (49). ZIKV NS4B interacts with SSR3 of the TRAP complex in neuroblastoma cells and ZIKV is dependent on at least two of the four TRAP complex subunits for infection of GSC, hiPSC-NPC and HeLa cells (Figure 5A). Further supporting our observation of ZIKV interacting and being dependent on the mammalian translocon is its dependence on SRPRB, SPCS3 and TRAM1; subunits of the Signal Recognition Particle (SRP), the Signal Peptidase Complex (SPCS) and the Translocating chain-associated membrane protein (TRAM), respectively. Interestingly, the viral protease NS2B-3 also interacts with subunits of the OST complex (STT3A, RPN1), Sec61 complex (SEC61B) and TRAP complex (SSR3) (Figure 6). These interactions likely facilitate the co-translational cleavage of the viral polypeptide by NS2B-3 into its individual viral proteins

Here, we identify that ZIKV NS4B and NS2B-3 directly interact with the core complexes of the mammalian translocon, and propose that these interactions are essential for the ZIKV life cycle in neuroblastoma cells. The dependency of ZIKV likely stems from the mammalian translocon facilitating viral polyprotein co-translational translocation, viral polyprotein cleavage, viral membrane protein insertion and/or viral protein N-glycosylation. Additionally, ZIKV may utilise its protein interactions with the Sec61 complex, TMEM33 and VAPB, to regulate the IRE1- and PERK-mediated UPR ER stress responses, that we identify to be significantly upregulated at the transcriptome level in ZIKV infected-neuroblastoma cells.

## SUMMARY

Our meta-analysis highlights the strong therapeutic potential of ZIKV, specifically the PRVABC59 strain, against multiple neuroblastoma cell-lines. We identify ZIKV to interact with, and be dependent on, multiple host protein complexes and pathways for its life cycle in paediatric neuroblastoma cells and for inducing oncolysis (Figure 7). Although this area of research is still at an early stage, our extensive survey of neuroblastoma ZIKV infection studies clearly demonstrates the potential of a ZIKV-based therapeutic. There are a few avenues which need to be addressed to progress this area of research, including; (1) assessing ZIKV’s oncolytic effect against neuroblastoma in xenograft mouse models, (2) assessing ZIKV’s capability to induce an anti-tumoral immune response against neuroblastoma in immune-competent *in vivo* models, and (3) considering the effectiveness and safety of employing different forms of ZIKV-based therapeutics against neuroblastoma. Examples of the latter may include live attenuated ZIKV strains or the construction of a virotherapy that collectively expresses ZIKV NS4B and CCL2, which we show here to hold elements of ZIKV’s oncolytic and immune activatory potential, respectively.

**Figure 7.**
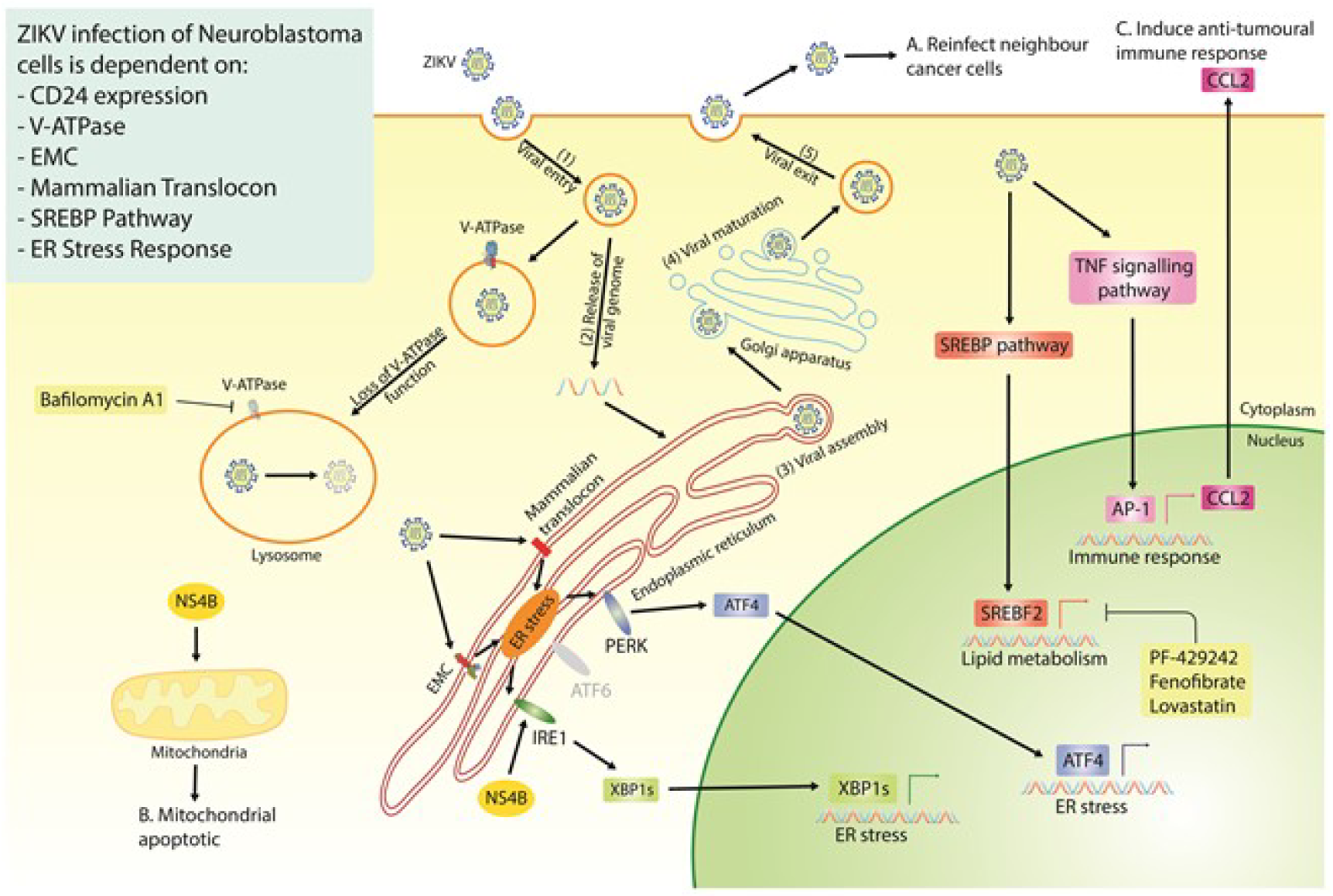
Diagram of the proposed ZIKV life cycle in neuroblastoma cells (Step 1-5), with a summary of all the currently known dependencies that the virus has for infection of neuroblastoma cells. Highlighted are the three essential properties of an oncolytic virus; the production of fresh viral particles to infect additional cancer cells (A), the ability to induce cancer cell death (B) and a mechanism through which ZIKV may induce an anti-tumoral immune response (C).

## ACKNOWLEDGEMENTS

The authors would like to thank the original curators of the datasets used in this study. Immense thanks goes to the various project funders, whose support made this work possible. These funders are (in order of funds received): MRC, Rosetrees Trust, Wessex Medical Trust, Little Princess Trust and Neuroblastoma UK.

## REFERENCES

1. Johnsen JI, Dyberg C, Wickström M. Neuroblastoma—A Neural Crest Derived Embryonal Malignancy. Front Mol Neurosci. 2019;12:9.

2. Campos Cogo S, Gradowski Farias da Costa do Nascimento T, de Almeida Brehm Pinhatti F, de França Junior N, Santos Rodrigues B, Cavalli LR, et al. An overview of neuroblastoma cell lineage phenotypes and in vitro models. Exp Biol Med Maywood NJ. 2020 Dec;245(18):1637– 47.

3. Irwin MS, Naranjo A, Zhang FF, Cohn SL, London WB, Gastier-Foster JM, et al. Revised Neuroblastoma Risk Classification System: A Report From the Children’s Oncology Group. J Clin Oncol [Internet]. 2021 Jul 28 [cited 2021 Dec 13]; Available from: https://ascopubs.org/doi/pdf/10.1200/JCO.21.00278

4. Chung C, Boterberg T, Lucas J, Panoff J, Valteau-Couanet D, Hero B, et al. Neuroblastoma. Pediatr Blood Cancer. 2021;68(S2):e28473.

5. Macedo N, Miller DM, Haq R, Kaufman HL. Clinical landscape of oncolytic virus research in 2020. J Immunother Cancer. 2020 Oct 1;8(2):e001486.

6. Li C, Xu D, Ye Q, Hong S, Jiang Y, Liu X, et al. Zika Virus Disrupts Neural Progenitor Development and Leads to Microcephaly in Mice. Cell Stem Cell. 2016 Jul 7;19(1):120–6.

7. Tang H, Hammack C, Ogden SC, Wen Z, Qian X, Li Y, et al. Zika Virus Infects Human Cortical Neural Precursors and Attenuates Their Growth. Cell Stem Cell. 2016 May 5;18(5):587–90.

8. Musso D, Ko AI, Baud D. Zika Virus Infection — After the Pandemic. Longo DL, editor. N Engl J Med. 2019 Oct 10;381(15):1444–57.

9. Adachi K, Nielsen-Saines K. Zika Clinical Updates: Implications for Pediatrics. Curr Opin Pediatr. 2018 Feb;30(1):105–16.

10. Zhu Z, Mesci P, Bernatchez JA, Gimple RC, Wang X, Schafer ST, et al. Zika Virus Targets Glioblastoma Stem Cells through a SOX2-Integrin αvβ5 Axis. Cell Stem Cell. 2020 Feb;26(2):187–204.e10.

11. Kaid C, Goulart E, Caires-Junior LC, Araujo BHS, Soares-Schanoski A, Bueno HMS, et al. Zika Virus Selectively Kills Aggressive Human Embryonal CNS Tumor Cells In Vitro and In Vivo. Cancer Res. 2018 Jun 15;78(12):3363–74.

12. Kaid C, Madi RA dos S, Astray R, Goulart E, Caires-Junior LC, Mitsugi TG, et al. Safety, Tumor Reduction, and Clinical Impact of Zika Virus Injection in Dogs with Advanced-Stage Brain Tumors. Mol Ther. 2020 May;28(5):1276–86.

13. Mazar J, Li Y, Rosado A, Phelan P, Kedarinath K, Parks GD, et al. Zika virus as an oncolytic treatment of human neuroblastoma cells requires CD24. PloS One. 2018;13(7):e0200358.

14. Scaturro P, Stukalov A, Haas DA, Cortese M, Draganova K, Płaszczyca A, et al. An orthogonal proteomic survey uncovers novel Zika virus host factors. Nature. 2018 Sep;561(7722):253–7.

15. Wang S, Zhang Q, Tiwari SK, Lichinchi G, Yau EH, Hui H, et al. Integrin αvβ5 Internalizes Zika Virus during Neural Stem Cells Infection and Provides a Promising Target for Antiviral Therapy. Cell Rep. 2020 Jan;30(4):969–983.e4.

16. Li Y, Muffat J, Javed AO, Keys HR, Lungjangwa T, Bosch I, et al. Genome-wide CRISPR screen for Zika virus resistance in human neural cells. Proc Natl Acad Sci. 2019 May 7;116(19):9527–32.

17. Savidis G, McDougall WM, Meraner P, Perreira JM, Portmann JM, Trincucci G, et al. Identification of Zika Virus and Dengue Virus Dependency Factors using Functional Genomics. Cell Rep. 2016 Jun 28;16(1):232–46.

18. Pereira R, Costa V, Gomes G, Campana P, Pádua R, Barbosa M, et al. Anti-Zika virus activity of plant extracts containing polyphenols and triterpenes on Vero CCL-81 and human neuroblastoma SH-SY5Y cells. Chem Biodivers. 2022 Mar 13;

19. Kedarinath K, Fox CR, Crowgey E, Mazar J, Phelan P, Westmoreland TJ, et al. CD24 Expression Dampens the Basal Antiviral State in Human Neuroblastoma Cells and Enhances Permissivity to Zika Virus Infection. Viruses. 2022 Aug 6;14(8):1735.

20. Anfasa F, Siegers JY, van der Kroeg M, Mumtaz N, Stalin Raj V, de Vrij FMS, et al. Phenotypic Differences between Asian and African Lineage Zika Viruses in Human Neural Progenitor Cells. mSphere. 2017 Aug;2(4).

21. Jorgačevski J, Korva M, Potokar M, Lisjak M, Avšič-Županc T, Zorec R. ZIKV Strains Differentially Affect Survival of Human Fetal Astrocytes versus Neurons and Traffic of ZIKV-Laden Endocytotic Compartments. Sci Rep [Internet]. 2019 May 30 [cited 2020 May 2];9. Available from: https://www.ncbi.nlm.nih.gov/pmc/articles/PMC6542792/

22. Bonenfant G, Meng R, Shotwell C, Badu P, Payne AF, Ciota AT, et al. Asian Zika Virus Isolate Significantly Changes the Transcriptional Profile and Alternative RNA Splicing Events in a Neuroblastoma Cell Line. Viruses. 2020 May 5;12(5):E510.

23. Hu H, Zhang W, Huang D, Wang Y, Zhang Y, Yi Y, et al. Clinical characteristics, treatment and prognosis of paediatric patients with metastatic neuroblastoma to the brain. Clin Neurol Neurosurg. 2019 Sep 1;184:105372.

24. Wen F, Armstrong N, Hou W, Cruz-Cosme R, Obwolo LA, Ishizuka K, et al. Zika virus increases mind bomb 1 levels, causing degradation of pericentriolar material 1 (PCM1) and dispersion of PCM1-containing granules from the centrosome. J Biol Chem. 2019 Dec 6;294(49):18742–55.

25. Gazon H, Barbeau B, Mesnard JM, Peloponese JM. Hijacking of the AP-1 Signaling Pathway during Development of ATL. Front Microbiol [Internet]. 2018 [cited 2022 Mar 29];8. Available from: https://www.frontiersin.org/article/10.3389/fmicb.2017.02686

26. Atsaves V, Leventaki V, Rassidakis GZ, Claret FX. AP-1 Transcription Factors as Regulators of Immune Responses in Cancer. Cancers. 2019 Jul 23;11(7):E1037.

27. Lima MC, Mendonça LR de, Rezende AM, Carrera RM, Aníbal-Silva CE, Demers M, et al. The Transcriptional and Protein Profile From Human Infected Neuroprogenitor Cells Is Strongly Correlated to Zika Virus Microcephaly Cytokines Phenotype Evidencing a Persistent Inflammation in the CNS. Front Immunol [Internet]. 2019 [cited 2020 Jul 9];10. Available from: https://www.frontiersin.org/articles/10.3389/fimmu.2019.01928/full

28. Braz-De-Melo HA, Pasquarelli-do-Nascimento G, Corrêa R, das Neves Almeida R, de Oliveira Santos I, Prado PS, et al. Potential neuroprotective and anti-inflammatory effects provided by omega-3 (DHA) against Zika virus infection in human SH-SY5Y cells. Sci Rep [Internet]. 2019 Dec 27 [cited 2020 May 2];9. Available from: https://www.ncbi.nlm.nih.gov/pmc/articles/PMC6984748/

29. Parker JN, Meleth S, Hughes KB, Gillespie GY, Whitley RJ, Markert JM. Enhanced inhibition of syngeneic murine tumors by combinatorial therapy with genetically engineered HSV-1 expressing CCL2 and IL-12. Cancer Gene Ther. 2005 Apr;12(4):359–68.

30. Osuna-Ramos JF, Reyes-Ruiz JM, del Ángel RM. The Role of Host Cholesterol During Flavivirus Infection. Front Cell Infect Microbiol. 2018 Nov 2;8:388.

31. Raini SK, Takamatsu Y, Dumre SP, Urata S, Mizukami S, Moi ML, et al. The novel therapeutic target and inhibitory effects of PF-429242 against Zika virus infection. Antiviral Res. 2021 Aug;192:105121.

32. Weber LW, Boll M, Stampfl A. Maintaining cholesterol homeostasis: Sterol regulatory element-binding proteins. World J Gastroenterol WJG. 2004 Nov 1;10(21):3081–7.

33. Sabino C, Basic M, Bender D, Elgner F, Himmelsbach K, Hildt E. Bafilomycin A1 and U18666A Efficiently Impair ZIKV Infection. Viruses. 2019 Jun 6;11(6).

34. Mlera L, Offerdahl DK, Dorward DW, Carmody A, Chiramel AI, Best SM, et al. The liver X receptor agonist LXR 623 restricts flavivirus replication. Emerg Microbes Infect. 2021 Jun 24;1–33.

35. Gladwyn-Ng I, Cordón-Barris L, Alfano C, Creppe C, Couderc T, Morelli G, et al. Stress-induced unfolded protein response contributes to Zika virus–associated microcephaly. Nat Neurosci. 2018 Jan;21(1):63–71.

36. Tan Z, Zhang W, Sun J, Fu Z, Ke X, Zheng C, et al. ZIKV infection activates the IRE1-XBP1 and ATF6 pathways of unfolded protein response in neural cells. J Neuroinflammation [Internet]. 2018 Sep 21 [cited 2020 Apr 30];15. Available from: https://www.ncbi.nlm.nih.gov/pmc/articles/PMC6151056/

37. Carr M, Gonzalez G, Martinelli A, Wastika CE, Ito K, Orba Y, et al. Upregulated expression of the antioxidant sestrin 2 identified by transcriptomic analysis of Japanese encephalitis virus-infected SH-SY5Y neuroblastoma cells. Virus Genes. 2019 Oct 1;55(5):630–42.

38. Bonenfant G, Meng R, Shotwell C, Badu P, Payne AF, Ciota AT, et al. Asian Zika Virus Isolate Significantly Changes the Transcriptional Profile and Alternative RNA Splicing Events in a Neuroblastoma Cell Line. Viruses. 2020 May;12(5):510.

39. Ngo AM, Shurtleff MJ, Popova KD, Kulsuptrakul J, Weissman JS, Puschnik AS. The ER membrane protein complex is required to ensure correct topology and stable expression of flavivirus polyproteins. eLife. 8:e48469.

40. Barrows NJ, Anglero-Rodriguez Y, Kim B, Jamison SF, Le Sommer C, McGee CE, et al. Dual roles for the ER membrane protein complex in flavivirus infection: viral entry and protein biogenesis. Sci Rep. 2019 Jul 4;9(1):9711.

41. Lin DL, Inoue T, Chen YJ, Chang A, Tsai B, Tai AW. The ER Membrane Protein Complex Promotes Biogenesis of Dengue and Zika Virus Non-structural Multi-pass Transmembrane Proteins to Support Infection. Cell Rep. 2019 May;27(6):1666–1674.e4.

42. Mangieri LR, Mader BJ, Thomas CE, Taylor CA, Luker AM, Tse TE, et al. ATP6V0C Knockdown in Neuroblastoma Cells Alters Autophagy-Lysosome Pathway Function and Metabolism of Proteins that Accumulate in Neurodegenerative Disease. PLOS ONE. 2014 Apr 2;9(4):e93257.

43. Li M, Zhang D, Li C, Zheng Z, Fu M, Ni F, et al. Characterization of Zika Virus Endocytic Pathways in Human Glioblastoma Cells. Front Microbiol [Internet]. 2020 Mar 6 [cited 2020 Jun 4];11. Available from: https://www.ncbi.nlm.nih.gov/pmc/articles/PMC7069030/

44. Owczarek K, Chykunova Y, Jassoy C, Maksym B, Rajfur Z, Pyrc K. Zika virus: mapping and reprogramming the entry. Cell Commun Signal. 2019 May 3;17(1):41.

45. Han X, Wang J, Yang Y, Qu S, Wan F, Zhang Z, et al. Zika Virus Infection Induced Apoptosis by Modulating the Recruitment and Activation of Proapoptotic Protein Bax. J Virol [Internet]. 2021 Mar 25 [cited 2021 Apr 7];95(8). Available from: https://jvi.asm.org/content/95/8/e01445-20

46. Braunger K, Pfeffer S, Shrimal S, Gilmore R, Berninghausen O, Mandon EC, et al. Structural basis for coupling of protein transport and N-glycosylation at the mammalian endoplasmic reticulum. Science. 2018 Apr 13;360(6385):215–9.

47. Puschnik AS, Marceau CD, Ooi YS, Majzoub K, Rinis N, Contessa JN, et al. A small molecule oligosaccharyltransferase inhibitor with pan-flaviviral activity. Cell Rep. 2017 Dec 12;21(11):3032–9.

48. Gwon YD, Zusinaite E, Merits A, Överby AK, Evander M. N-glycosylation in the Pre-Membrane Protein Is Essential for the Zika Virus Life Cycle. Viruses. 2020 Aug 23;12(9):E925.

49. Monel B, Compton AA, Bruel T, Amraoui S, Burlaud-Gaillard J, Roy N, et al. Zika virus induces massive cytoplasmic vacuolization and paraptosis-like death in infected cells. EMBO J. 2017 Jun 14;36(12):1653–68.

50. Hughes BW, Addanki KC, Sriskanda AN, McLean E, Bagasra O. Infectivity of Immature Neurons to Zika Virus: A Link to Congenital Zika Syndrome. EBioMedicine. 2016 Jun 23;10:65–70.

51. Bagasra O, Shamabadi NS, Pandey P, Desoky A, McLean E. Differential expression of miRNAs in a human developing neuronal cell line chronically infected with Zika virus. Libyan J Med. 2021 Jan 1;16(1):1909902.

52. Mlera L, Bloom ME. Differential Zika Virus Infection of Testicular Cell Lines. Viruses [Internet]. 2019 Jan 9 [cited 2020 Apr 30];11(1). Available from: https://www.ncbi.nlm.nih.gov/pmc/articles/PMC6356326/

53. Hou W, Armstrong N, Obwolo LA, Thomas M, Pang X, Jones KS, et al. Determination of the Cell Permissiveness Spectrum, Mode of RNA Replication, and RNA-Protein Interaction of Zika Virus. BMC Infect Dis [Internet]. 2017 Mar 31 [cited 2020 May 1];17. Available from: https://www.ncbi.nlm.nih.gov/pmc/articles/PMC5374689/

54. Castro FL, Geddes VEV, Monteiro FLL, Gonçalves RMDT, Campanati L, Pezzuto P, et al. MicroRNAs 145 and 148a Are Upregulated During Congenital Zika Virus Infection. ASN Neuro. 2019 Jan;11:175909141985098.

55. Mendonça-Vieira LR de, Aníbal-Silva CE, Menezes-Neto A, Azevedo E de AN, Zanluqui NG, Peron JPS, et al. Reactive Oxygen Species (ROS) Are Not a Key Determinant for Zika Virus-Induced Apoptosis in SH-SY5Y Neuroblastoma Cells. Viruses. 2021 Oct 20;13(11):2111.

56. Bos S, Viranaicken W, Turpin J, El-Kalamouni C, Roche M, Krejbich-Trotot P, et al. The structural proteins of epidemic and historical strains of Zika virus differ in their ability to initiate viral infection in human host cells. Virology. 2018 Mar 1;516:265–73.

57. Sánchez-San Martín C, Li T, Bouquet J, Streithorst J, Yu G, Paranjpe A, et al. Differentiation enhances Zika virus infection of neuronal brain cells. Sci Rep [Internet]. 2018 Sep 28 [cited 2020 Apr 30];8. Available from: https://www.ncbi.nlm.nih.gov/pmc/articles/PMC6162312/

58. Giel-Moloney M, Goncalvez AP, Catalan J, Lecouturier V, Girerd-Chambaz Y, Diaz F, et al. Chimeric yellow fever 17D-Zika virus (ChimeriVax-Zika) as a live-attenuated Zika virus vaccine. Sci Rep [Internet]. 2018 Sep 4 [cited 2020 Apr 30];8. Available from: https://www.ncbi.nlm.nih.gov/pmc/articles/PMC6123396/

59. Haviernik J, Štefánik M, Fojtíková M, Kali S, Tordo N, Rudolf I, et al. Arbidol (Umifenovir): A Broad-Spectrum Antiviral Drug That Inhibits Medically Important Arthropod-Borne Flaviviruses. Viruses [Internet]. 2018 Apr 10 [cited 2020 May 2];10(4). Available from: https://www.ncbi.nlm.nih.gov/pmc/articles/PMC5923478/

60. Alpuche-Lazcano SP, McCullogh CR, Del Corpo O, Rance E, Scarborough RJ, Mouland AJ, et al. Higher Cytopathic Effects of a Zika Virus Brazilian Isolate from Bahia Compared to a Canadian-Imported Thai Strain. Viruses [Internet]. 2018 Jan 27 [cited 2020 Apr 30];10(2). Available from: https://www.ncbi.nlm.nih.gov/pmc/articles/PMC5850360/

61. Himmelsbach K, Hildt E. Identification of various cell culture models for the study of Zika virus. World J Virol. 2018 Feb 12;7(1):10–20.

62. Offerdahl DK, Dorward DW, Hansen BT, Bloom ME. Cytoarchitecture of Zika virus infection in human neuroblastoma and Aedes albopictus cell lines. Virology. 2017 Jan;501:54–62.

